# DeePNAP: A deep learning method to predict protein-nucleic acids binding affinity from sequence

**DOI:** 10.1101/2023.12.03.569768

**Authors:** Uddeshya Pandey, Sasi M. Behara, Siddhant Sharma, Rachit S. Patil, Souparnika Nambiar, Debasish Koner, Hussain Bhukya

## Abstract

Predicting the protein-nucleic acid (PNA) binding affinity solely from their sequences is of paramount importance for the experimental design and analysis of PNA interactions (PNAIs). A large number of currently developed models for binding affinity prediction are limited to specific PNAIs, while also relying on both sequence and structural information of the PNA complexes for both train/test and also as inputs. As PNA complex structures available are scarce, this significantly limits the diversity and generalizability due to a small training dataset. Additionally, a majority of the tools predict a single parameter such as binding affinity or free energy changes upon mutations, rendering a model less versatile for usage. Hence, we propose DeePNAP, a machine learning-based model trained on a vast and heterogeneous dataset with 14,401 entries (from both eukaryotes and prokaryotes) of ProNAB database, consisting of wild-type and mutant PNA complex binding parameters. Our model precisely predicts the binding affinity and free energy changes due to the mutation(s) of PNAIs exclusively from the sequences. While other similar tools extract features from both sequence and structure information, DeePNAP employs sequence-based features to yield high correlation coefficients between the predicted and experimental values with low root mean squared errors for PNA complexes in predicting the *K*_D_ and ΔΔG implying the generalizability of DeePNAP. Additionally, we have also developed a web interface hosting DeePNAP that can serve as a powerful tool to rapidly predict binding affinities for a myriad of PNAIs with high precision toward developing a deeper understanding of their implications in various biological systems. Web interface: http://14.139.174.41:8080/

## Introduction

Protein-nucleic acids (PNA) binding affinity prediction based on their primary structural information can be of major interest as it readily indicates their interaction feasibility. PNA interactions (PNAIs) are indispensable in maintaining cellular physiology, responding to various environmental stimuli, host-pathogen interaction, and comprehension of the PNA binding affinity can provide an edge over manipulating these interactions to promote healthcare and fitness. These interactions frequently dictate the fate of cellular metabolism, signal transfer, and other processes to adapt to the recurring and often harsh environmental conditions through genetic reprogramming^1^. PNAIs, along with other biomolecular interactions, are primarily responsible to overcome stress and maintain homeostasis and are hence of paramount importance^2–4^. Knowledge of the interaction energy of PNA, and fine-tuning their binding affinity may eventually help to envision how these transient interactions operate in biological systems^5^. For example, mutations in DNA/RNA binding proteins leading to their dysfunction is at the root of many diseases^6–10^. Therefore, it is vital to perform the qualitative and quantitative characterization of PNAIs.

The current methods to achieve this qualitative and quantitative data are *via* interaction studies like gel shift assays, calorimetry, fluorescence-based techniques, nuclear magnetic resonance spectroscopy (NMR), etc. along with *in-silico* methods like molecular dynamics^11–14^. The three-dimensional (3D) structure elucidation of PNA complexes experimentally is a reliable method for identifying binding residues that result in interaction specificity. However, determining the 3D structure of PNA complexes using cryo-electron microscopy, X-ray crystallography, NMR, and other experimental approaches is laborious, time-consuming, and require proteins to be isolated in high quantities and purity which often yields a lower success rate^15^. As a result, only 13,234 entries to date (November, 2023) are available when searching for protein-DNA/RNA complex structures in public databases like Protein Data Bank (PDB)^16^. Moreover, estimating the binding affinity due to the interactions is equally laborious and may eventually arrive at a similar low success rate scenario. Computational PNA docking is another method for creating 3D structures of the complex which require individual structures of protein and cognate NA. Other *in-silico* approaches for calculating the binding affinity of PNA rely on molecular dynamics simulations to aid *in-silico* docking and virtual screening^17–22^. Accuracy and computational resources are seldom mutually exclusive^23^. Hence, binding affinity determination or estimation of various PNA complexes is one of the long-standing challenges in experimental and computational biology.

Alternatively, this challenge can be addressed by making use of the currently available experimental PNAI profiles and quantitative binding affinity data on wild and mutant PNA complexes. With this comprehensive data in hand, there is a pressing need to convert it into meaningful information using recent developments in artificial intelligence (AI) and machine learning (ML)^24–25^. The ability of the ML model to “learn” intrinsic patterns in a complex plane of available data has resulted in resource-optimal predictions that do not compromise accuracy. Employing these technological advancements, PNAI strength can be quantified rapidly from their sequence, especially when the existing experimental procedures fall short. A few examples where sequence information is extensively used as an input for an ML-based model include AlphaFold2^26^, OmegaFold^27^, and RoseTTAFold^28^ which are extensively used in 3D structure prediction with outstanding performance, and DeepTFactor used to predict transcription factors^29^.

A number of currently available ML based, as well as other models predict the binding affinities or free energy changes for PNA complexes; however, most of them suffer from two major drawbacks: (1) the usage of structural data to train, test and prediction and (2) a significantly small dataset used for training and testing. Additionally, a number of models are limited to measure a specific type of PNA interactions, such as protein-DNA or protein-RNA interactions. Some such examples include the PreDBA and PredPRBA, which are ML-based heterogeneous ensemble and gradient boosted regression tree models, respectively, that require structure as input and are trained on a relatively small dataset of 100 protein-DNA and 103 protein-RNA complexes, respectively.^30,31^ Another ML-based tool, SAMPDI-3D which is a gradient boosting decision tree machine learning method, was trained on a total of 883 entries with 419 protein and 463 DNA mutants and also uses structural data as input.^32^ Other tools such as mCSM-NA and mmCSM-NA, which relies on graph-based structural signatures and emPDBA which is an ensemble regression model were trained on 331 (including mutants), 155 experimentally solved complexes and 340 entries respectively.^33–35^ The dataset contained approximately 700 P-DNA complexes, however, this was refined in order to train and test DNAffinity, while PDA-Pred was trained on 391 entries. Both of these models are ML-based approaches and derives features from structural and/or simulation data.^36,37^ Apart from ML-based tools, PremPDI uses optimized parameters from experimental sets of mutations (219 in total, with 49 unique protein-DNA complexes).^38^ As observed, most of the above-mentioned tools extract features from both structure and sequences of PNA complexes to train their models on a relatively small dataset. This may result in the lack of adequate descriptors that can account for the poor description of the complexity in binding partners. Moreover, using structural data for model training introduces a tight bottleneck towards method accuracy and generalizability, which occurs due to the significant challenges faced during the structural elucidation of proteins themselves, let alone PNA complexes. Thus, there is a pressing need to develop a reliable and versatile prediction tool that can capture the diverse nature of PNAIs using a large training dataset and also functions using sequence information alone.

In order to develop a model with high prediction accuracy and enable generalizability over diverse PNAIs, a reliable and efficient deep neural network-based ML model, DeePNAP is reported in this work. DeepNAP estimates the binding affinity (order of dissociation constant) and free energy changes (upon mutations in PNA) exclusively from sequence information only. It is trained and tested on a dataset of 14,401 entries (5,881 unique entries) as given in the ProNAB^39^ database representing a relatively large sample size. Each entry further yields sequence-derived information such as polar, charged and hydrophobic nature of individual residues and also the information on the partner NA sequence. The model was further subjected to hundred-fold random data splitting to assess the model performance, which resulted in an average Pearson correlation (R) value of between 0.86, an average root mean squared error (RMSE) of 0.83 and an average mean absolute error (MAE) of 0.57 in predicting *K*_D_ for wild and mutant PNA complexes. An ensemble model using five sets of optimized DeePNAP parameters was also developed in this work for both wild and mutant PNA complexes. The ensemble model has an R, RMSE and MAE values of 0.93, 0.63 and 0.44, respectively, in predicting *K*_D_ for both wild and mutant PNA complexes. This indicates the generalizability of DeePNAP which is commendable considering the vast heterogeneity in the dataset curated from the ProNAB database. In comparison to the above-mentioned predicting tools, DeePNAP has shown an overall R-value of 0.96 for the entire dataset under study. Furthermore, DeePNAP has been trained to predict both *K*_D_ and ΔΔG independently enabling the users to draw informed conclusions, and has also shown comparable performance against currently available ΔΔG prediction tools. Hence, DeePNAP can serve as a powerful tool to predict binding affinities for a myriad of PNAIs with high precision, which can aid in the rapid development of a deeper understanding of the various implications of PNAIs in biological systems. DeePNAP allows the user to ascertain the prediction quality by comparing the thermodynamic binding parameter (ΔΔG and *K*_D_) predictions for a given complex, and has also been made available to predict the binding affinity and free energy changes upon mutations of PNA complexes *via* a user-friendly tool (Web interface: http://14.139.174.41:8080/. Additionally, the code to install DeePNAP locally is also made available at GitHub: https://github.com/StructuralBiologyLabIISERTirupati).

## MATERIAL AND METHODS

### Data processing

The deep neural network model developed in this work is named ‘DeePNAP’ and was trained and optimized on the ProNAB dataset^39^. Protein and the corresponding target NA sequences, their dissociation constants, and other thermodynamic binding parameters are tabulated in the ProNAB dataset. In total, more than 20,000 entries of protein-DNA and protein-RNA complexes, obtained from different experiments, are reported in this dataset. To construct a ML model predicting the binding affinity of PNA complexes, the dataset entries with proteins and NA sequences to a maximum length of 1,000 amino acids and 75 bases, respectively, were chosen. This work excluded several entries containing special characters/spaces, resulting in 14,401 total entries, with 11,071 wild and 3,330 mutant variants of PNA complexes considered in the model training and test datasets. A brief summary of the dataset is shown in **Figure 1** and **Figure S1**. Among the 14,401 entries, repetition found for several entries as the same PNA complexes have been studied and reported in many different references. For this purpose, we have used the average value of the orders of *K_D_*s for the same PNA complexes. After removing the repeated entries, the dataset contains 5,881 entries of wild and mutant PNA complexes. Out of 5,881 entries, there are 4,397 wild PNA complexes and 1,484 mutant PNA complexes.

**Figure 1.**
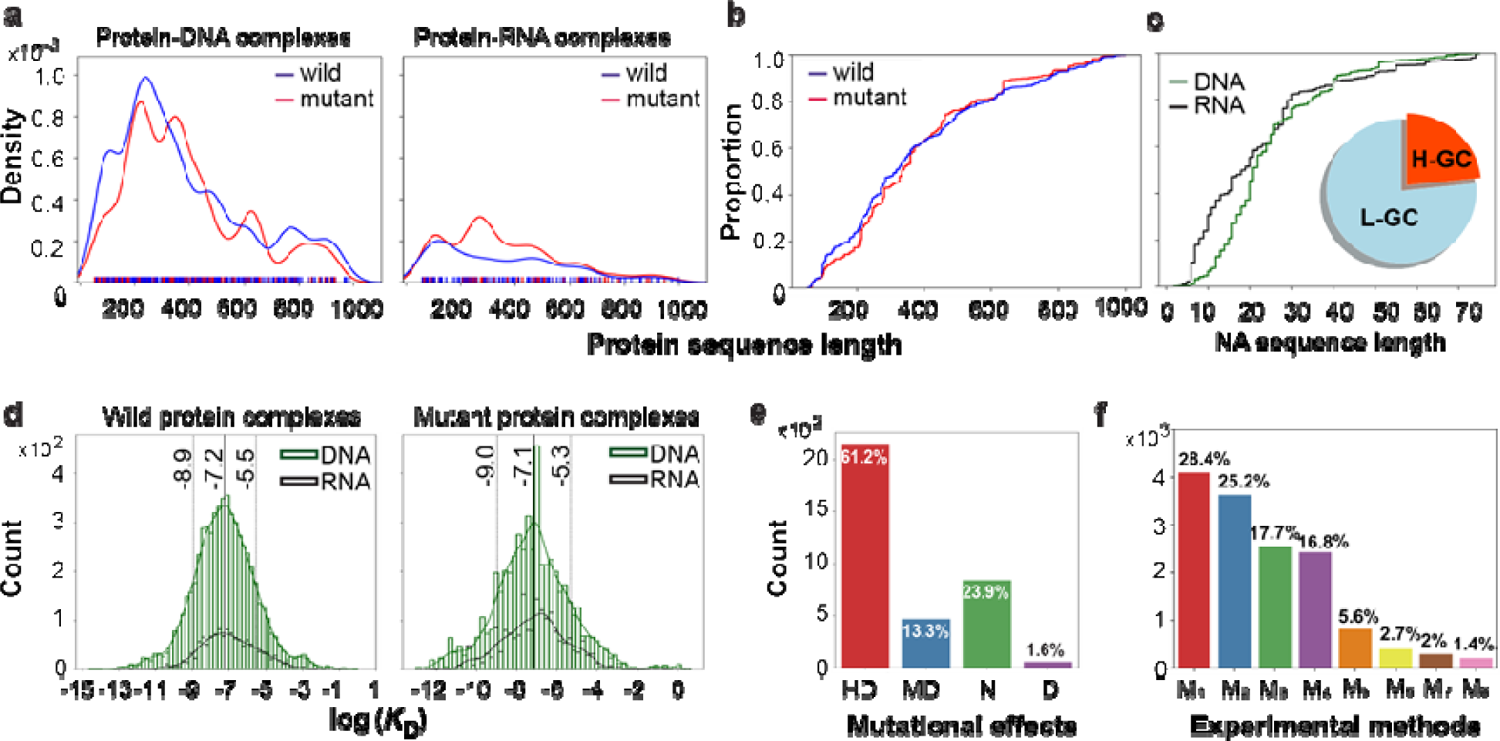
Filtered dataset statistics. a) Distribution of wild and mutant proteins shown in blue and red solid lines, respectively, for protein sequences in the PNA complexes. b,c) Indicate the cumulativ frequency reaching saturation at the protein (wild and mutant) and NA sequence lengths of 1,000 and 75, respectively, in the PNA complexes. A pie chart in the inset shows the proportion of NA sequences with high (H-GC) and low (L-GC) GC content. Sequences containing more than 60% GC content are considered H-GC. d) Distribution of dissociation constants (*K*_D_) of wild and mutant proteins in the PNA complexes. The normal distributions of protein-DNA and protein-RNA complexes are represented as solid green and black curves, respectively, for both wild and mutant PNA complexes. The solid and dotted lines indicate the mean and standard deviation, respectively, of the distribution. e) The bar graph displays the effect of mutations (HD, highly destabilizing; MD, moderately destabilizing; N, neutral; and D, destabilizing) in the protein on the changes in the free energy change (ΔΔG). f) Contribution of the entries from broadly classified experimental methods used to measure the *K*_D_ of protein-NA binding complexes in the filtered dataset. Where each technique is represented with a bar plot. Gel shift assays (M1), fluorescence (M2), calorimetry (M3), filter binding measurements (M4), surface plasmon resonance (M5), quenching (M6), other (M7) experiments, and footprinting assays (M8).

### Descriptors and Encoding

Proteins and nucleic acids are expressed in terms of the amino acid and nucleobase symbols, respectively, which are typically distinct letters or objects. Thus, a protein or nucleic acid is a sequence of characters. An embedding technique is needed to convert those discrete objects to numerical values to be used as descriptors in an ML model. One-hot encoding of each amino acid or nucleobase is the most common choice for datasets related to the PNA complex, which represents a protein or nucleic acid with a 2D sparse matrix. However, different algorithms can be used in order to reduce the dimensionality of the problem. In this work, a combination-based encoding algorithm in which a vector with 6 elements representing each amino acid was employed. Each of those elements can have a value of either ‘0’ or ‘1’ and each amino acid is represented by a vector with three ‘1’ and three ‘0’. Thus, all 20 possible combinations represent 20 amino acids. For example, lysine is represented as ‘[1, 0, 0, 1, 0, 1]’. Since the embedding is performed randomly, all amino acids belong to different categories. However, amino acids can be divided into different categories based on the side chain’s physical properties like polarity, hydrophobicity, and charge. In order to incorporate the physical properties of the side chain of amino acids, a three-element vector was augmented to the already embedded six-element vector for each amino acid. The augmented vectors for amino acids with charged, polar, and hydrophobic side chains are [1, 0, 0], [0, 1, 0], and [0, 0, 1], respectively. Thus, an amino acid is represented by a nine-element vector, e.g., lysine is represented as [1, 0, 0, 1, 0, 1, 1, 0, 0]. Using the embedding mentioned above, a protein sequence is encoded in an embedding matrix of dimension *N_p_*×9 (*N_p_* = the number of amino acid residues in the protein). A maximum chain length of 1,000 residues was considered for this work. For proteins shorter than 1,000 amino acids, the embedding matrix was padded with zeros to make the dimension of the protein descriptor 1,000×9. NA sequences are embedded using a one-hot encoding for five nucleobases. For example, adenine was encoded as [1, 0, 0, 0, 0]. The maximum number of nucleobases considered in this work is 75 which represents the nucleobase descriptor as a 75×5 matrix. For shorter nucleic acids (less than 75 bases), the embedding matrix was padded with zeros. Thus, for the ML model, each sample (protein-nucleic acid pair) is represented by two 2D descriptors, one for protein and another for NA.

### Model architecture

Our ML model consists of three modules (i) feature extraction module (FEM), (ii) interaction module (IM), and (iii) prediction module (PM). The FEM extracts the features from the input descriptors and passes the important messages to the IM. In the IM, the messages interact between themselves and finally send refined information to the PM to predict the desired output, see **Figure 2**. The architecture of the modules is discussed below.

**Figure 2.**
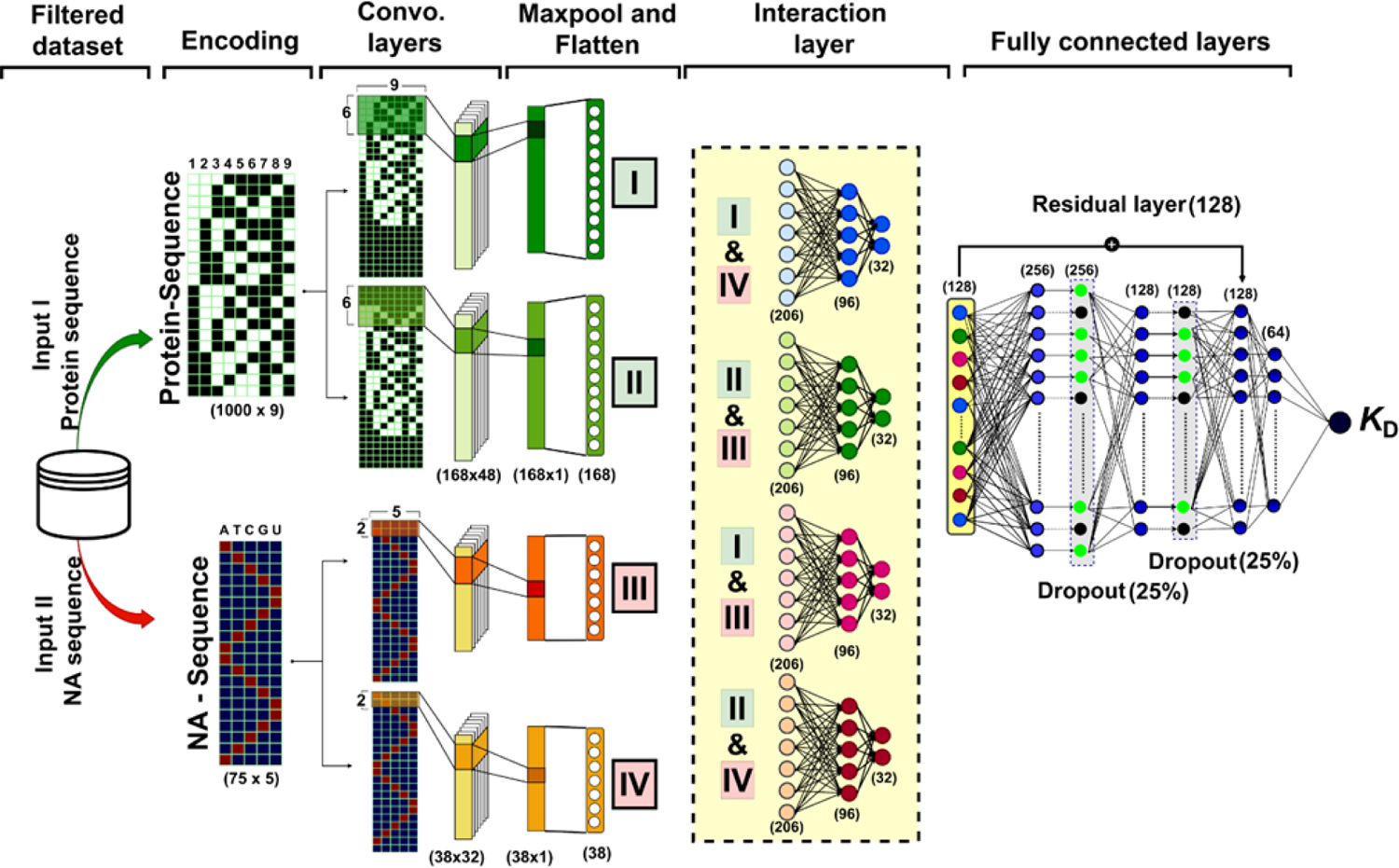
The architecture of the ML model, DeePNAP developed in this work. The input protein sequence from the filtered dataset is encoded into a matrix (1000×9) using combinatorial (features: 1-6) and physical property (features: 7-9) parameters (see text for details). One-hot encoding embeds the NA sequence into a (75×5) matrix. The features of these embedded matrices are extracted via a pair of parallel convolutional layers with different receptive fields from each protein and NA sequence. The important features extracted from the PNA complex via the convolutional layers are used in the interaction layers. Each of the four possible combinations of the extracted features was passed through a different neural network. Subsequently, the outcome from the interaction layers is fed to a set of fully connected dense layers to estimate the binding affinity, *K*_D_ or its changes, ΔΔG for the given PNA complex.

#### Feature extraction module (FEM)

In our ML model, we constructed a feature extraction module with four branches consisting of convolution layers and pooling layers to extract the important features from the encoded protein and NA sequences in one-dimensional (1D) representations. In FEM, two branches extract the features from the protein descriptors, while the other two branches find out the features from the NA descriptors. Each branch of the FEM consists of (i) a padding layer, (ii) a convolutional layer, (iii) a pooling layer, and (iv) a flattening layer. The padding layer is used to match the dimensionality for the next layer and to shift the receptive field of convolution in a customized way. A set of 48 filters of dimension 6×9 were used in the convolution layer with a stride of 6 along the chain length. After the convolution layer, a max pooling layer is added, which pools the maximum features across all the channels. Finally, the pooled features are flattened to obtain a 1D output. The two branches that extract the protein features are similar except that for one branch, eight dummy residues (with all zeros) were padded at the end of the encoded sequences, and for another branch, 3 dummy residues were padded at the beginning of the sequence so that the filter operates in a 3-unit shifted fashion (see **Figure 2** and **Figure S2**).

The basic architecture of the layers for the branch that extracts features from the nucleic acid sequences is similar to the ones used for the protein. A set of 32 filters of dimension 2×5 was used in the convolution layer with a stride of 2 along the chain length. In one branch, one dummy residue was padded at the end of the encoded sequences, whereas in another branch, one dummy residue was padded at the beginning of the sequence, so that the filter operates in a one-unit shifted fashion. By using stride, the whole protein/NA sequences were divided into small blocks and the convolution operation was applied to each block separately. Each individual block was the receptive field for a single convolution operation. In the next step, for another branch, the receptive field was shifted by half of the filter length along the direction of the chain length, this ensures some connectivity between two adjacent blocks.

#### Interaction module (IM)

In the interaction module, the four 1D output vectors obtained from the FEM interact with each other *via* a feed-forward neural network. The IM has four parallel blocks. Each block consists of a fully connected neural network containing two dense layers of 96 and 32 neurons. The inputs for the four blocks in IM are the four different combinations of the four outputs generated by the four branches (B1-4) of FEM (two outputs (I, II) from B1 and B2 branches and the other two outputs (III, IV) from B3 and B4 branches). Thus, the inputs for the four blocks of IM are the 1D sequences obtained *via* concatenation of the pairs (I, III), (I, IV), (II, III), and (II, IV). Dropout^40^ of 50% and L1 regularization were applied to the first and second dense layers, respectively, see **Figure 2**.

#### Prediction module (PM)

In this module, the log_10_(*K*_D_) of a protein-NA pair is predicted using the outputs generated by the IM. PM is a feed-forward neural network with four fully connected dense layers with 256, 128, 64, and 1 neurons. The input for this module is a concatenated array of the four 1D output vectors obtained from the four blocks of IM. In this module, a shortcut connection was used by adding the input to the output of the second dense layer. Fifty percent dropout was used for the first two layers, and L1 regularization was used in the third layer. Finally, the output was obtained from the last layer with a single neuron, see **Figure 2**. The python code snippet of the model is given in the SI as **Scheme 1** for reference.

### Model optimization

Keras API^41^ was used in this work to build and train the ML model. As mentioned before, after excluding the incomplete/unusable entries and repetition a total of 5,881 protein and NA pairs (4,397 wild and 1,484 mutant proteins) were used to optimize the ML model. In most of the mutant proteins, the difference between the wild and mutant sequences was minor (differs by one or two units). Since the ML model only uses sequence information as input, the difference between the predicted binding affinities for wild and mutant proteins and NA is very small. This leads to a small ΔΔ*G* value predicted by the ML model for many samples. To overcome this problem, in the DeepNAP model, two parallel neural networks (NN) of the same architecture as DeePNAP were used. One NN was predicting the log_10_(*K^w^*) for the wild protein and NA pair, while another one was predicting the log_10_(*K^m^*_D_) for the mutant protein and NA pair. The architecture of the ML model is shown in **Figure S3**. The loss function contains information regarding the change of binding affinities upon mutation, and thus the ML model learns the correlation between the wild and mutant proteins. This strategy significantly improved the prediction for the mutant protein and NA pair. Finally, the ML model was trained several times, as shown in **Figure S3**, and the five best sets of parameters were saved. The final model is an ensemble model that uses these five sets of parameters and predicts the average value obtained using each set of parameters separately.

### Baseline methods to compare with DeePNAP performance

In order to compare the performance of DeepNAP, we have used several baseline methods e.g., (i) decision tree (DT), with a maximum depth of 10, (ii) multiple linear regression (MLR), (iii) support vector machine (SVM), and (iv) random forest (RF), with 10 trees and a maximum depth of 10. The scikit-learn package^42^ was used in this work to optimize the baseline methods. The default values were used for the hyperparameters of the models unless mentioned otherwise. The sequences for proteins and NAs are encoded as one-dimensional vectors by using integer encoding for the basic units (amino acids or nucleobases) to feed them into the models.

### DeePNAP performance in comparison with existing tools

To perform comparative analysis of DeePNAP performance with mmCSM-NA, mCSM-NA, SAMPDI-3D and PremPDI in predicting the ΔΔG, the PremPDI dataset was adopted.^36^ The dataset has 219 mutant entries from 49 unique protein-DNA complex structures. Out of these 219 entries, the mmCSM-NA, SAMPDI-3D and mCSM-NA share 181, 77 and 105 entries with PremPDI. However, the entire DeePNAP’s test and train datasets have only 25 entries common with the PremPDI dataset owing for its performance on a relatively blind dataset.

## Results and Discussion

### Database refinement

Towards building a precise ML-based PNAI binding affinity prediction model exclusively from the sequence information, the ProNAB database was employed.^39^ A detailed statistical analysis of all 20,009 entries extracted from the ProNAB database (as of January, 2022) was performed to evaluate the data distribution. This was performed to see the variations in protein and NA sequence lengths, the number of wild and mutant entries with their thermodynamic binding parameters, experimental methods for binding affinity measurements, and the type of PNA (DNA/RNA) complex. The summary statistics of the ProNAB data set are presented in **Figure 1** and **Figure S1**. The saturation was observed for protein and NA sequences at lengths of 1,000 residues and 75 bases, respectively, based on their respective cumulative frequency distributions, **Figure S1**. Additionally, any modified sequences with special characters were also eliminated to reduce statistically insignificant entries in the dataset. On execution of the above operations, 14,401 entries were sieved in the filtered dataset, with 11,244 protein-DNA and 3,157 protein-RNA complexes, which consisted of 11,071 wild-type protein and 3,330 mutant protein complexes, out of which 5,881 entries are unique in the dataset. The dataset was randomly split into training and test sets in a ratio of 9:1. The variation/conservation in the resultant protein and NA sequences was assessed to probe for model biasness introduced due to skewed training data, which showed minimal conservation/high variance. The training and test datasets together exhibited 6.8% and 13.1% identity for protein and nucleic acid sequences, while the individual datasets showed 6.5% and 12.6% identities for training, and 6.9% and 13.1% identities for test dataset in protein and nucleic acids sequences, respectively, see **Figure S4**. Protein sequence lengths for both wild (blue) and mutant (red) PNA complexes show approximately normal distribution, as represented in **Figure 1a**.

The saturation in the cumulative frequency distribution of protein and NA sequence lengths prompted their capping to retain maximum data variations in the filtered dataset while minimizing computation power, see Figure 1b,c and **Figure S1**. Further, we analyzed the binding affinity trends in the filtered dataset, which showed a normal distribution for wild and mutant PNA complexes, with the mean observed for the log_10_(*K*_D_) at −7.2 ± 1.7 M and −7.1 ± 1.8 M respectively, Figure 1d. Additionally, to assess the quality of mutant complexes in the filtered dataset, the distribution of changes in free energy changes (ΔΔG) was also plotted, Figure 1e. The ΔΔG (kcal/mol) values –0.25 ≤ ΔΔG ≤ +0.25, –2.00 ≤ ΔΔG < −0.25, –4.00 ≤ ΔΔG < −2.00, and –4.00 < ΔΔG are categorized into neutral (N), mildly destabilizing (MD), destabilizing (D), and highly destabilizing (HD) according to the report.^43^ From Figure 1e, it was observed that the ΔΔG values were mainly HD (∼60%) and N (∼23%) mutational effects. The experimental binding affinity and ΔΔG measurements were obtained from various techniques, as shown in Figure 1f. The results of this plot suggest that the majority of the filtered dataset entries are from gel shift assays (M1) followed by fluorescence (M2), calorimetry (M3), and filter binding measurements. However, there are also a significant number of entries for which the binding affinity and ΔΔG estimations were obtained by performing surface plasmon resonance (M5), quenching (M6), footprinting assays (M8), and other (M7) experiments. This non-homogenous distribution in the entries obtained from different experimental methods employed (as mentioned above) to determine the binding affinity and the free energy changes has its own intrinsic errors in the measurements. These errors will eventually be fed to the model in order to train and test it, and as a result, it is highly unlikely for any model to reach maximum prediction accuracy. Due to the inevitability of such errors in other similar tools, it is necessary to develop a robust model that can effectively minimize these sources of error and reliably predict binding affinity parameters for uncharacterized PNAIs. This has been achieved for DeePNAP with the use of a large and diverse curated dataset consisting of broad sequence length and binding affinity distributions, mutational effects, and PNAI strength determination methods. Thus, DeePNAP can potentially represent a tool that can precisely predict thermodynamic binding parameters for a spectrum of PNAIs.

### DeePNAP performance

The refined dataset used in this work contains wild- and mutant-type proteins and NA complexes which were used to train the DeePNAP model (architecture and feature extraction module shown in Figure 2 and **Figure S2** respectively).

The DeepNAP model has been trained 100 times with a training *vs* test split of 9:1 using random splitting of the dataset. For each of the splitting, the model’s trainable parameters were initialized 4 times. Here it is worth mentioning that for each of the runs, the test set was never used during training. Also, early stop criteria were used to prevent overfitting. The results obtained from those 100 runs are summarized in Figure 3. Compared to the experimental values, the predictions of the optimized models have an average R of 0.96 for the training datasets and 0.86 for the validation datasets. The average RMSE values in predicting *K*_D_ for PNA complexes are 0.49 and 0.83, respectively for the training and validation datasets. The RMSEs for the mutant PNA complexes are a bit higher than the wild PNA complexes and also the predictions for the mutant PNAIs have a higher standard deviation. The average values of R for the ΔΔG are 0.92 and 0.6, for training and test sets respectively. This substantial difference signifies that the optimized model somewhat lacks interpreting the binding sites of PNA complexes. However, it should be noted that there is also scarcity of data availability at present for mutant PNA complexes and nevertheless the model generalization ability can be increased with the availability of more data in future. The R, RMSE and MAE values obtained from the best model are 0.92, 0.67 and 0.48 respectively in predicting *K*_D_, while the R, RMSE and MAE values are 0.77, 1.04 kcal/mol, and 0.74 kcal/mol respectively for ΔΔG.

**Figure 3.**
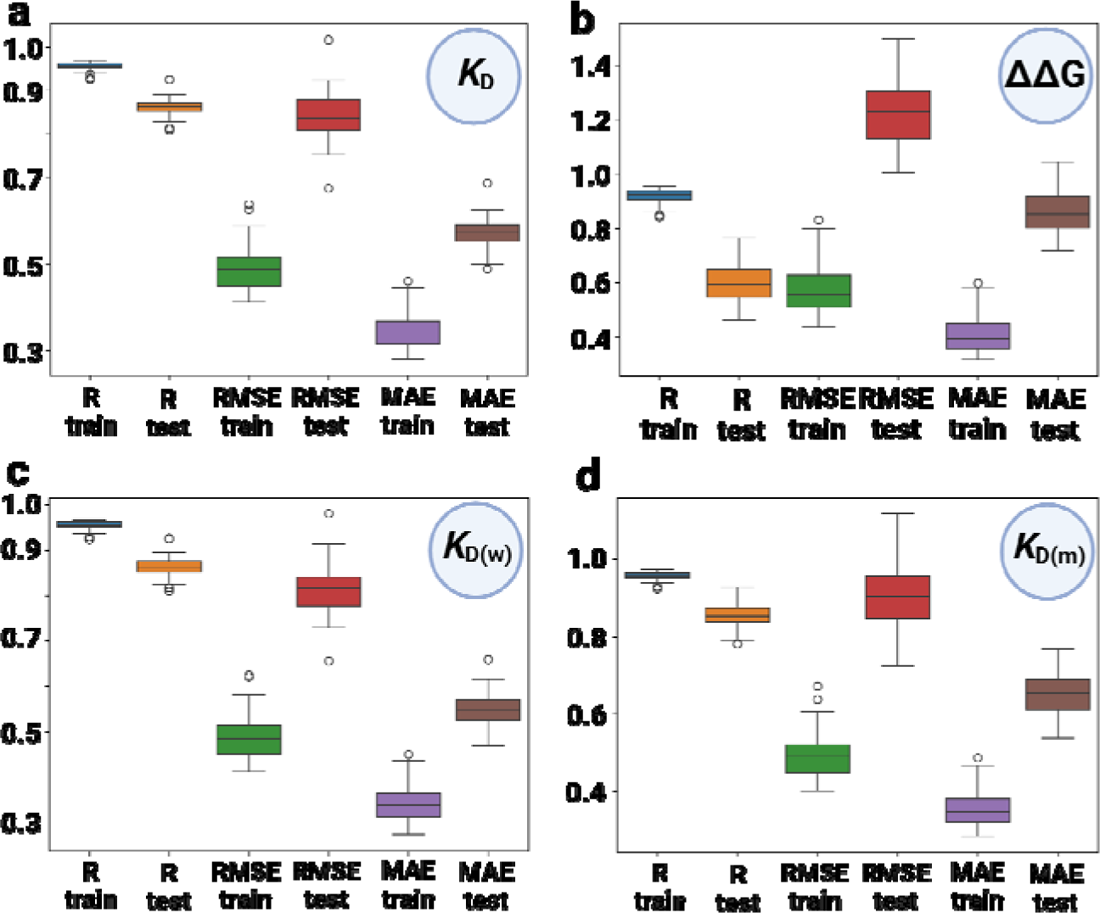
Benchmarking the model performance by 100 random splitting of the data in a 9:1 ratio of training vs test sets. a) *K*_D_ correlation values for all (wild and mutant) protein and NA pairs along with root mean squared error (RMSE) and mean absolute error (MAE) for test and train sets. b) ΔΔG correlation values along with RMSE and MAE for test and train sets. *K*_D_ correlation values for all wild (c) and mutant (d) protein and NA pairs along with RMSE and MAE for test and train sets. The unit of *K*_D_ i the order (10^n^) while the unit of ΔΔG is kcal/mol.

The best model was retrained for several rounds, and an ensemble model was constructed using the five best sets of parameters. The performance of the optimized model is shown in Figure 4.

**Figure 4.**
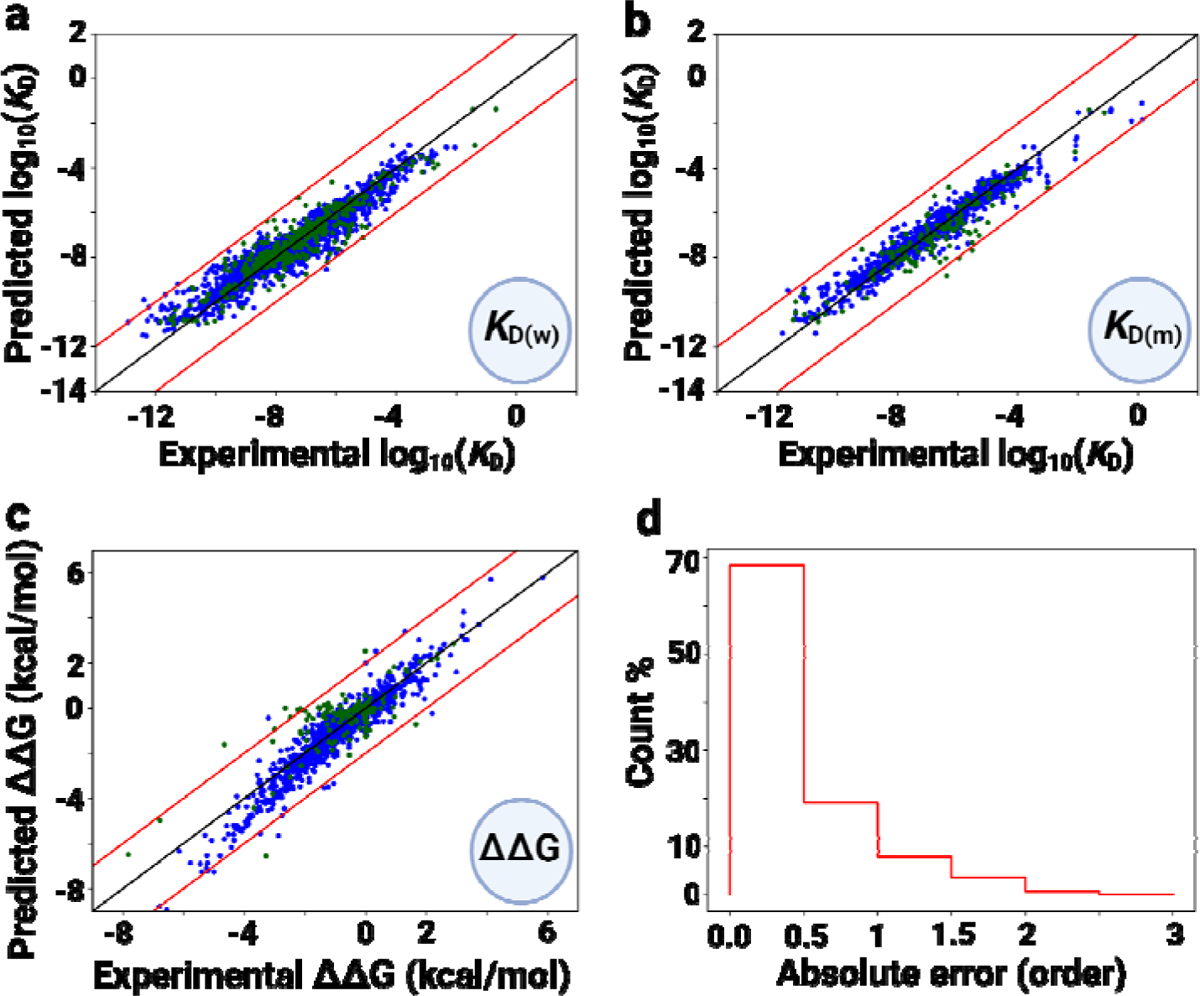
DeePNAP performance on the final dataset containing wild and mutant proteins, and NA pairs. a) Correlation between experimental and predicted log_10_(*K*_D_) values for wild protein and NA pairs. b) Correlation between experimental and predicted log_10_(*K*_D_) values for mutant proteins and NA pairs. c) Correlation between experimental and predicted ΔΔG values for protein-NA interaction upon mutation of the protein. In panels (a,b,c), the blue-filled circles represent the samples used in training while the olive filled circles represent the samples used for validation purposes. The black diagonal lines in panels (a,b,c) show a perfect correlation while the red diagonal lines show a deviation of ±2 compared to the experimental values. d) Histogram plot of the sample count with respect to the absolute error in predicting experimental log_10_(*K*_D_) values for the validation set. The X-axis in panel (d) is the absolute error in predicting the order (10^n^) of the dissociation constant for PNA complexes.

The prediction accuracy for PNA binding affinity over wild and mutant complexes is shown in Figure 4a,b. The R values for the wild and mutant entries from the training set are 0.96 and 0.97, respectively, whereas for the validation set, R values are 0.94 and 0.93 for the wild and mutant PNA complexes, respectively. The RMSE values for the wild and mutant samples were 0.43 and 0.39, respectively, for the training set, while the E values for the wild and mutant samples were 0.59 and 0.74, respectively, for the validation set. From the result, it was evident that the model is efficient in yielding predictions that are in close agreement with the experimentally derived binding affinities. Additionally, DeePNAP was also tested for its ability to estimate changes in free energy (ΔΔG) upon mutation(s) in the NA binding proteins. The results are shown in Figure 4c which shows a good correlation to predict ΔΔG from only the sequence information of the protein and NA.

The R values for the data reported in Figure 4c are 0.95 and 0.81 for the training and validation sets, respectively. The mean absolute errors in predicting ΔΔG were 0.54 and 0.97 kcal/mol for the training and validation data, respectively. Interestingly, it was observed that the deviations in the binding affinity predictions for both wild and mutant protein NA complexes lie within two orders of magnitude, while the majority (about ∼85%) of the predicted *K*_D_ remain within one order as experimentally derived affinities (see Figure 4d). Thus, DeePNAP (see **Figure S6** for the input page of the web interface) is able to indicate the effects of the mutation(s) on various PNAIs.

### Comparison with baseline methods

The baseline methods (DT, MLR, SVM and RF) were run for several times and the best results are tabulated in **Table 1** along with the DeepNAP model. While the RF model performs best among the baseline models, DeepNAP’s predictions are significantly better than all baseline methods. It is also found that the MLR method fails to predict the binding affinity from sequence only inputs.

**Table 1.** Comparison of DeepNAP performance with baseline methods.

| Models | R($K_d$) | RMSE ($K_d$) | MAE ($K_d$) | R($\Delta\Delta G$) | RMSE($\Delta\Delta G$) (kcal/mol) | MAE($\Delta\Delta G$) (kcal mol) |
| --- | --- | --- | --- | --- | --- | --- |
| DeepNAP | **0.93** | **0.67** | **0.49** | **0.77** | **1.04** | **0.74** |
| DT | 0.86 | 0.90 | 0.63 | 0.48 | 1.61 | 1.0 |
| MLR | 0.007 | 131696 | 8625 | -0.048 | 458730 | 22492 |
| SVM | 0.82 | 1.05 | 0.69 | 0.42 | 3.5 | 1.37 |
| RF | 0.91 | 0.77 | 0.57 | 0.59 | 1.32 | 0.98 |

### Sensitivity of DeePNAP performance on its architecture

Since the proteins and NAs are encoded in 2D matrices here, the convolution layers are included in the beginning of the model to extract the important features and map it into one-dimensional vectors. 2D encoding of protein and NA is needed when only the sequence information is fed to a model. A similar architecture was also used in DeepTFactor^29^. The sensitivity of different modules of DeepNAP on the prediction of binding affinity is checked by comparing with modified versions of DeepNAP. A modified version referred to as ‘1’, the prediction module was truncated by using only one dense layer with a single neuron. Another modified version was also tested, where the B2 and B4 blocks were removed from the feature extraction module. This leads to only one block in the interaction module. This version is referred to as ‘2’. The original DeepNAP model is referred to as ‘0’. The R, RMSE and MAE obtained by running these three versions of the models 50 times are shown in the **Figure S5**. It is quite obvious that removing or truncating a particular module significantly reduces the model performance.

### DeePNAP performance comparison with existing prediction methods

A number of prediction models to quantify PNAIs have been developed earlier; however, their versatility and accuracy remains limited. For instance, PreDBA and PredPRBA were able to achieve averaged Pearson correlation coefficients (R) 0.84 and 0.8, with 0.87 and 1.11 as mean absolute errors respectively. SAMPDI-3D, on the other hand, was able to achieve a Pearson’s correlation coefficient of 0.73 and 0.43 with mean squared errors of 0.72 and 0.90 for cross-validation and blind test, respectively. PDApred achieved a correlation of 0.86 and a mean absolute error of 0.76 in self-consistency. mCSM-NA and emPDBA achieved a correlation of 0.68 and 0.53 on blind tests. In contrast, mmCSM-NA achieved a correlation of 0.67 (with a RMSE of 1.06 kcal/mol) when evaluating single-point mutations through cross-validation. Additionally, it demonstrated a correlation of up to 0.65 on independent non-redundant datasets involving multiple-point mutations (with a root mean square error of 1.12 kcal/mol). PremPDI achieved a correlation of 0.71 on its training and testing dataset, with root mean squared error of 0.86. On the other hand, DNAffinity achieved a determination coefficient of 0.93 ± 0.02 on its training data from the gcPBM dataset, 0.69 ± 0.17 from uPBM dataset and 0.7 ± 0.14 on the HT-SELEX dataset. These numbers suggest the quality of the tool’s performances in predicting the free energy changes for the P-NA complexes on a relatively small dataset with the available 3D structures. This has become the bottleneck for developing an effective tool given the difficulties in solving the structures of the PNA complexes. Use of sophisticated techniques like electron cryo-microscopy to determine protein-NA complex structures is a way forward; however, the complex structures available as of today are subminimal^44,45^. On the other hand, there is a vast amount of experimental data available for the PNA interactions; their sequences are known with no available 3D structures. To overcome this, developing DeePNAP was an attempt to extract information from just the sequences and predict the unknown P-NA binding interactions.

In order to assess the efficiency of our developed model in comparison to other tools, we tested the performance of DeePNAP against four latest prediction tools, namely: mmCSM-NA, mCSM-NA, SAMPDI-3D and PremPDI. These tools were chosen for comparison in accordance to their compatibility with PremPDI dataset. It should also be noted that mCSM-NA and SAMPDI-3D have previously been evaluated using the PremPDI dataset. Upon analysis, it was found that DeePNAP performance was comparable to all these tools, showing a correlation of 0.51 between the predicted and the experimental ΔΔG values. While mmCSM-NA, mCSM-NA, PremPDI and SAMPDI-3D yielded correlation coefficients of 0.56, 0.57, 0.71 and 0.83, respectively. The parameter for prediction (ΔΔG) was chosen since DeePNAP is trained on a smaller dataset for ΔΔG prediction as compared to the *K*_D_ prediction, and hence is anticipated to show a lower accuracy for ΔΔG prediction as compared to *K*_D_. It is also worth noting that all these tools are specifically trained for ΔΔG prediction; however, DeePNAP performance is on par with these tools, which are specifically trained to predict *K*_D_ values of PNA complexes. Additionally, these models rely on both sequence as well as 3D structure of the complex, where interaction information is already embedded in the input. Moreover, these tools are limited to a specific PNAI *i.e* protein-DNA interactions, whereas DeePNAP is able to predict binding parameters for both P-DNA and P-RNA complexes. As a result, DeePNAP effectively addresses the problem of carrying out PNAI predictions in spite of the scarcity of solved PNA complex structures by reliably predicting binding affinity parameters using features extracted from sequence alone. Thus, DeePNAP can indeed serve as a tool to predict binding parameters for various PNA complexes and can also help to narrow down to potential binding partners for uncharacterized sequences.

We have also compared the performance of DeepNAP with PreDBA^30^, which predicts the *K_D_*. The same dataset which was used in PreDBA was used here. As mentioned in the article, PreDBA yields a Pearson correlation 0.84 only when they build separate models for separate protein-DNA complexes. However, when all data were used to train a single model, the performance was rather poor with a Pearson correlation of 0.17. DeepNAP provides a better Pearson correlation of 0.73 on the same dataset.

## Conclusion

We propose DeePNAP, an ML-based model that can be highly useful for predicting the binding affinities and free energy changes for various PNA complexes using features exclusively extracted from their sequences. This model was developed using the PNA complexes, which include both wild-type and mutant protein-DNA and protein-RNA complexes from both prokaryotes and eukaryotes (see **Figure S7**). As a result of this vast training set, DeePNAP is able to distinguish between protein-DNA and protein-RNA complexes, while predicting thermodynamic PNAI parameters (binding strength and free energy changes) for wild-type and mutant PNA complexes with high accuracies in no time. DeePNAP employs features extracted exclusively from sequence and shows a high correlation value 0.86 for wild-type proteins and 0.85 for mutant-protein and NA complexes with a RMSEs of 0.90 and 0.81 for the wild and mutant PNA complexes, respectively. The final ensemble model predicts the log_10_(*K*_D_) for wild/mutant proteins and NA complexes with an RMSE value of 0.63 and a mean absolute error of 0.44 for the validation set. The model also shows a fair correlation with the experimental data in predicting the ΔΔG. The high prediction accuracy coupled with its ability to predict both *K*_D_ and ΔΔG independently makes DeePNAP one of the accurate and robust sequence-based PNAI parameter prediction tools that can significantly accelerate the experimental characterization of novel PNA complexes. Hence, it is anticipated that DeePNAP can provide a great platform for rapid, feasible, and reliable prediction of binding parameters for a myriad of PNAIs. In addition, DeePNAP is also made available as an open-source tool with a free, user-friendly web interface for the bioscience community.

### ASSOCIATED CONTENT

#### Data Availability Statement

ProNAB dataset is available at https://web.iitm.ac.in/bioinfo2/pronab/, Web server: http://14.139.174.41:8080/ GitHub: https://github.com/StructuralBiologyLabIISERTirupati.

#### Supporting Information

The following files are available free of charge. ProNAB data statistics, DeePNAP architecture, FEM, random data splitting and model performance, sequence identity, ML model, web-interface, and data distribution (PDF)

#### Author Contributions

The project was conceived and developed by H.B., U.P. and D.K. Data curation and processing by U.P., R.S.P., S.N., S.S. D.K. and H.B. Model development and performance evaluation by D.K., U.P., S.M.B., S.S. and H.B. Web interface developed by S.M.B. Results analysis and presentation by D.K., H.B., U.P., S.S. and R.S.P. The manuscript was written H.B., D.K., R.S.P., and U.P. with inputs from other authors. Revisions are made by D.K., H.B., S.M.B., R.S.P., S.S. and S.N. All authors have given approval to the final version of the manuscript. Project was supervised by H.B. and D.K. ^†^U.P. and S.M.B contributed equally as first authors and ^#^S.S. and R.S.P contributed equally as second authors.

## ACKNOWLEDGMENT

H.B. thanks IISER Tirupati for the infrastructure. U.P., S.S., R.S.P., S.M.B. and S.N., thank IISER Tirupati for fellowship and infrastructure. M.M. Gromiha for providing the ProNAB dataset. We thank Amogh Desai for the discussion on the project. D.K. thanks IISER Tirupati, IIT Hyderabad, and DST INSPIRE for the computational and financial support. H.B. thank Param Brahma of the National Supercomputing Mission for providing access to computational resources. All authors thank Srikanth Varma, Ravi Chavali and Satish Jadav of IISER Tirupati for IT support.

## ABBREVIATIONS

PNA: protein-nucleic acid

PNAI: PNA interactions

NA: nucleic acid

NMR: nuclear magnetic resonance spectroscopy

PDB: protein data bank

ML: machine learning

AI: artificial intelligence

MSA: multiple sequence alignment

RMSE: root mean squared error

MAE: mean absolute error

3D: three-dimension.

