## Supplementary Information for "DeePNAP: A deep learning method to predict protein-nucleic acids binding affinity from sequence"

**Biological Resources:**

Pronab dataset preparation: A data set containing the experimentally derived binding affinity parameters (binding free energy, dissociation, and association constants) of protein and nucleic acid complexes and the sequences of the protein sequence and the binding DNA was prepared from the ProNab dataset[1]. Among the 22,009 entries in the ProNab dataset, a protein-nucleic acid binding complex was included in this study if the entry satisfies the following criteria: 1) The absence of non-standard amino acid (e.g., B, X, O, U, Z) in the protein sequence (non-standard amino acid cite), 2) The absence of non-standard or modified bases in the nucleic acids, 3) The protein sequences containing only the standard 20 amino acids were included, 4) The nucleic acid sequences containing the standard bases like A, T, C, G and U. 5) Nucleic acid sequences below 75 bases in length were selected. 6) Only those complexes were included in the training data for which criteria 1-5 were positive and were mapped to a non-zero dissociation constant ($K_D$). After processing, the reconstructed data set contained 14,401 entries of the protein-nucleic acid complex with the statistics seen in **Figure S1**.

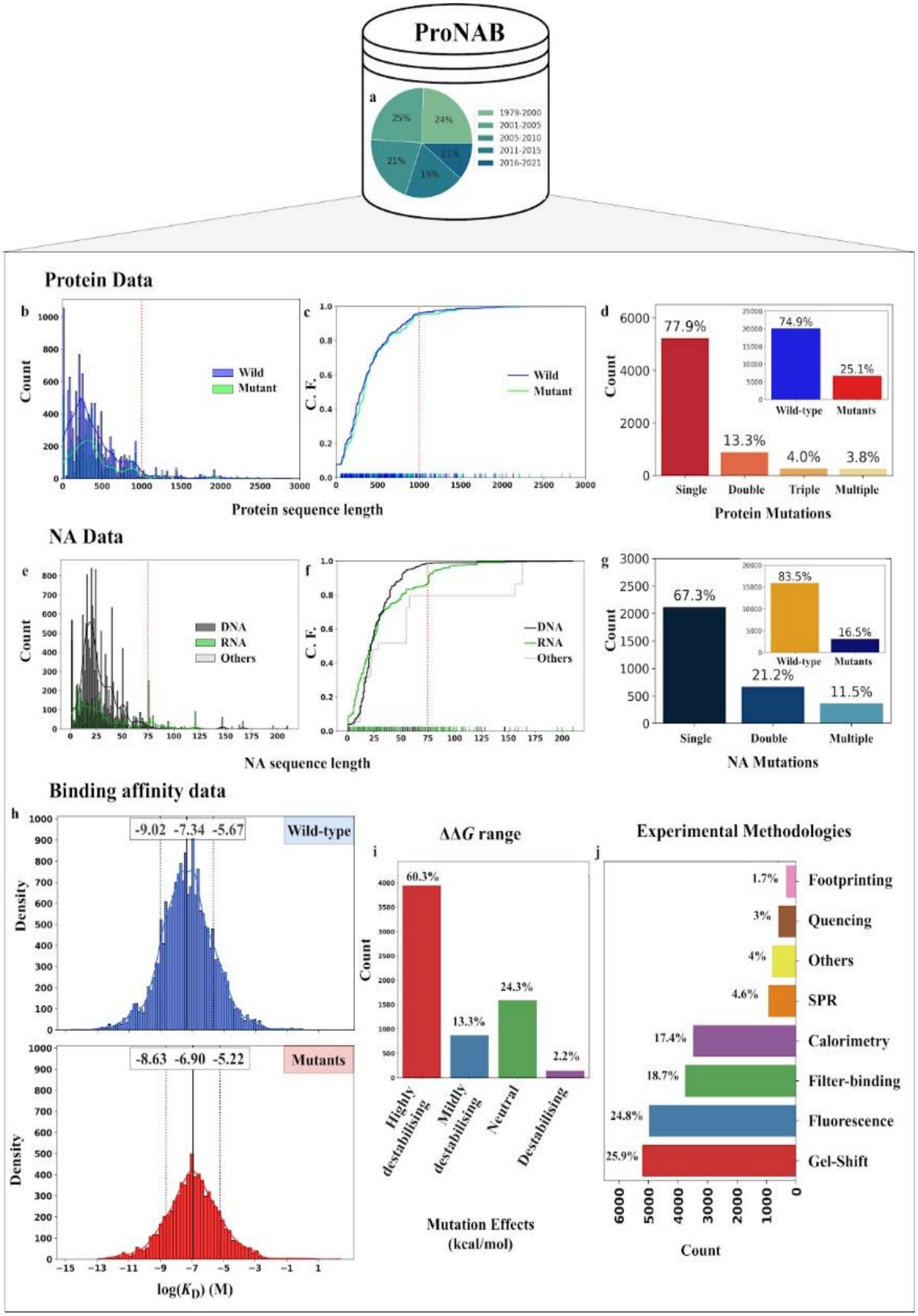


**Figure S1:** ProNAB dataset statistics. a) Year-wise distribution of entries of the dataset. b,c) Distribution of wild and mutant shown in blue and green solid lines, respectively, for protein sequences in the PNA complexes. The cumulative frequency reaching saturation at the protein (wild and mutant) sequence length of 1,000 in the PNA complexes are indicated with red dashed line. d) The bar graph shows the percentages of different mutations in the protein, and their percentage of wild and mutant entries in the ProNAB dataset. e,f) The cumulative frequency reaching saturation at the NA (wild and mutant) sequence length of 75 in the PNA complexes are indicated with red dashed line. g) The bar graph shows the percentages of different mutations in NA, and their percentage of wild and mutant entries in the dataset. h) Distribution of dissociation constants ($K_D$) of wild and mutant proteins in the PNA complexes. The normal distributions of wild-type and mutant PNA complexes are represented as solid blue and red curves, respectively. The solid and dotted lines indicate the mean and standard deviation, respectively, of the distribution. i) The bar graph displays the changes in the free energy change ($\Delta\Delta G$) upon mutations (HD, highly destabilizing; MD, moderately destabilizing; N, neutral; and D, destabilizing) in the protein. j) Contribution of the entries from broadly classified experimental methods used to measure the $K_D$ of protein-NA binding complexes in the filtered dataset. Where each technique is represented with a bar plot. Gel shift assays, fluorescence, calorimetry, filter binding measurements, surface plasmon resonance, quenching, footprinting assays, and other experiments.

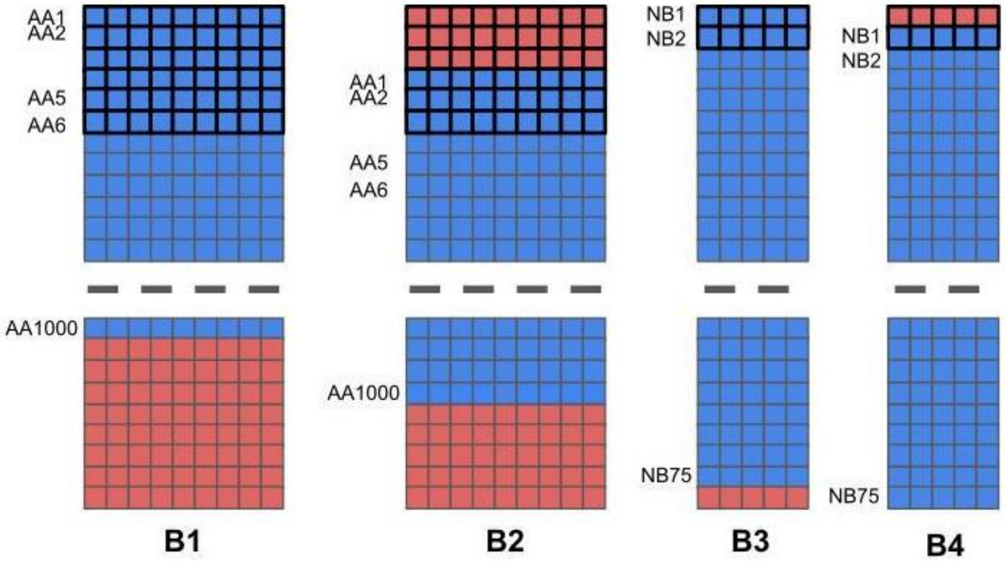


**Figure S2:** Padded input matrices and receptive fields for the four branches of FEM. B1 and B2 extract features from protein while B3 and B4 extract features from nucleic acids. The blue shaded blocks are the actual input matrix obtained after encoding, as described in the text. 'AA*i*' is *i*-th amino acid, while 'NB*i*' is *i*-th nucleobase. The red-shaded blocks are the zero padding regions, and the blocks with thick black borders show the receptive field for a single convolution operation.

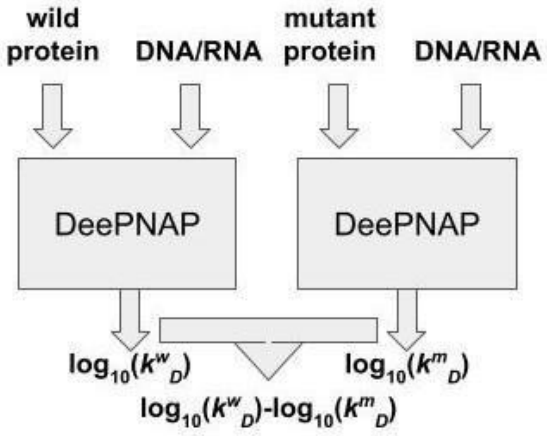


**Figure S3**. ML model to efficiently optimized DeePNAP to predict the binding affinity of wild-type proteins and NA pairs as well as mutant proteins and NA pairs. The ML model consisted of two DeePNAP models that shared the same set of weights. One model predicts the $\log_{10}(K^w{}_D)$ value for wild protein and NA pair, while the other predicts $\log_{10}(K^m{}_D)$ value for mutant protein and NA pair. The ML model has another output, the difference between both $\log_{10}(K_D)$ values, i.e., ($\log_{10}(K^w{}_D)$-$\log_{10}(K^m{}_D)$). The loss function is the sum of mean squared error for the three outputs weighted by 0.5, 0.5, and 1.5 for $\log_{10}(K^w{}_D)$, $\log_{10}(K^m{}_D)$, and the difference between them, respectively.

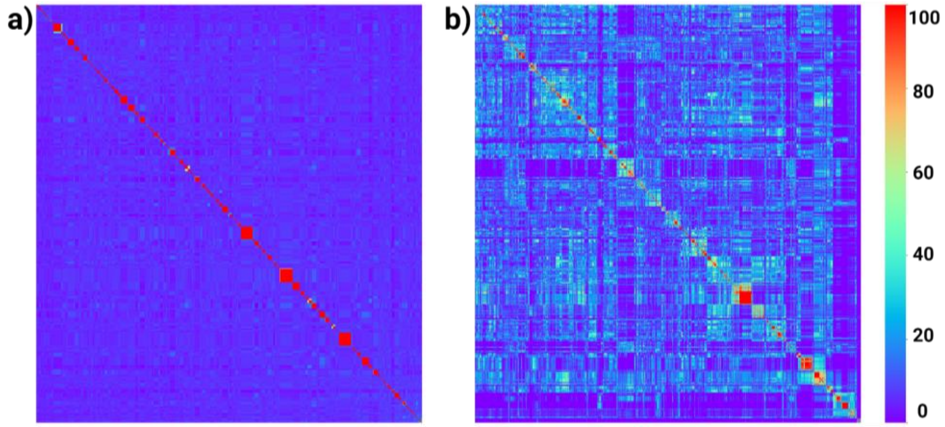


**Figure S4**. Heatmap for the pairwise sequence percentage identity among the combined test and train dataset. a) & b) Protein and nucleic acid sequence percentage identity, respectively. The color scale represents the level of sequence identity from 0% (violet) to 100% (red).

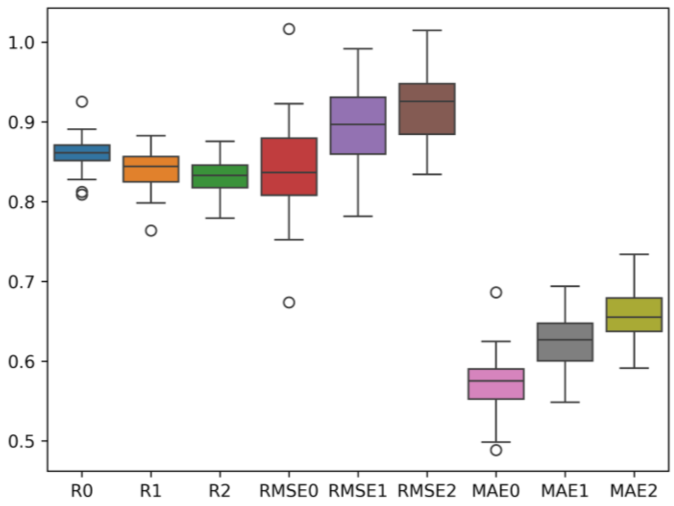


**Figure S5:** Performance comparison of different versions of DeepNAP model. The original DeepNAP model is referred as '0' while a modified version referred as '1', is generated by truncating the prediction module via using only one dense layer with a single neuron. In a second modified version, B2 and B4 blocks were removed from the feature extraction module. which leads to only one block in the interaction module. This version is referred as '2'. The Pearson correlation coefficients (Ri), root mean square errors (RMSEi) and mean absolute errors (MAEi) shown here are obtained by running the three versions of the models 50 times.

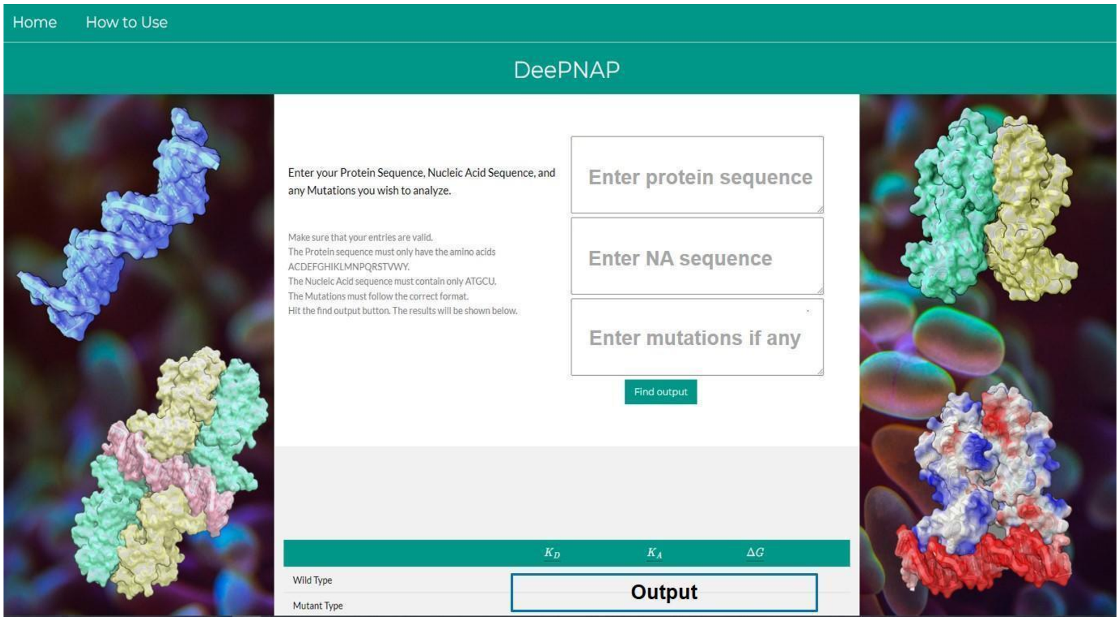


**Figure S6.** DeePNAP user interface showing the input page and the output being displayed at the bottom of this page.

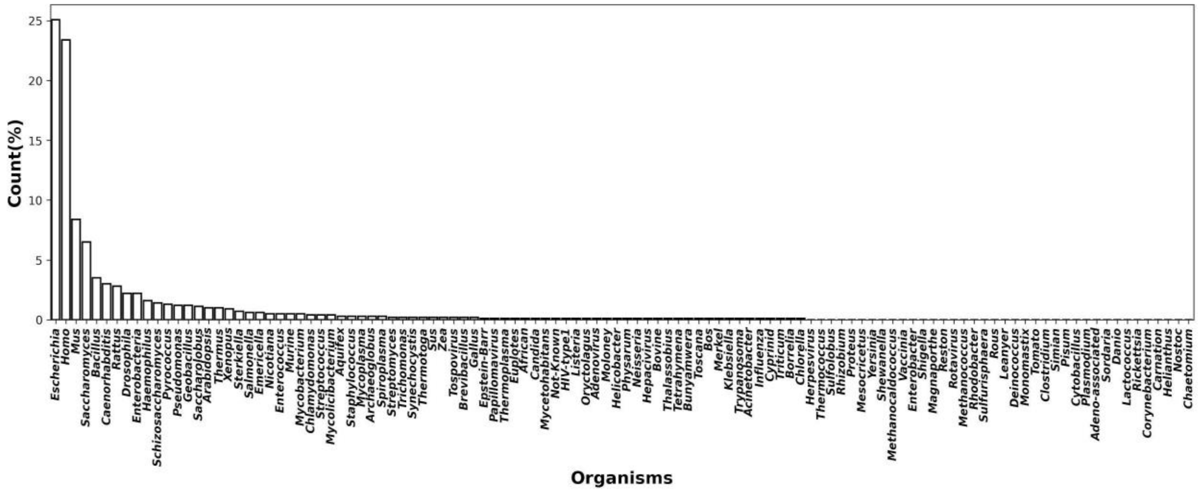


**Figure S7.** The percentile distribution of entries from various organisms in the filtered dataset is depicted in the histogram. The X-axis represents the genus names, while the corresponding percentage counts are visually displayed on the Y-axis. Organisms whose sources were not specified in the ProNAB dataset are labeled as 'Not-Known'.
